# A rapid, inexpensive, non-lethal method for detecting disseminated neoplasia in a bivalve

**DOI:** 10.1101/2023.06.28.544680

**Authors:** Lauren E. Vandepas, Ryan N. Crim, Emily Gilbertson, Marisa A. Yonemitsu, Elizabeth Unsell, Michael J. Metzger, Adam Lacy-Hulbert, Frederick W. Goetz

## Abstract

Disseminated neoplasia (DN) is a form of cancer in bivalve molluscs that has been reported in some cases to be a transmissible cancer. Neoplastic cells are highly proliferative, and infection is often lethal. Some commercially valuable bivalve species (mussels, cockles, soft-shell clams, oysters) are affected by outbreaks of disseminated neoplasia, making disease diagnosis and mitigation an important issue in aquaculture and ecological restoration efforts. Here we describe a minimally invasive, non-lethal method for high-throughput screening for disseminated neoplasia in basket cockles (*Clinocardium nuttallii*). Basket cockles are native to the North American Pacific coast from California to Alaska. There is recent concern from some Coast Salish Tribes regarding an observed long-term decline in cockle populations in Puget Sound, WA. This has led to increased interest in monitoring efforts and research to improve our understanding of the mechanisms of observed basket cockle population dynamics, including assessing prevalence of disease, such as disseminated neoplasia. The rapid, non-lethal hemolymph smear screening method presented here to diagnose DN in adult *C. nuttallii* can be applied at field sites at low financial cost, and in a validation study of 29 animals the results were identical to that of the gold standard method, tissue histology. Due to the similar morphology of DN in different bivalves, this method can likely be generally applied for use in any bivalve species.

## INTRODUCTION

Many bivalve species worldwide are susceptible to disseminated neoplasia (DN), a fatal disease in which cancerous cells proliferate in the hemolymph of the animal and eventually disseminate into solid tissues in later stages of disease (Barber 2004). Neoplastic cells are highly proliferative and most heavily localized to the hemolymph – the blood-like fluid in molluscs containing a mixture of nutrients, respiratory gases, and circulating cells – though the source cell-type of disseminated neoplastic cells has not yet been identified (Alderman et al.. 2017). Neoplastic cells migrate through unknown mechanisms to infiltrate other tissues in advanced stages of the disease (Barber et al. 2004; Alderman et al. 2017). Recently, DN in multiple bivalve species has been shown to be transmissible between individuals or, in some cases, between species (Metzger et al. 2015, Murchison 2016, Metzger et al. 2016, Yonemitsu et al. 2019). DN has been reported in diverse bivalve taxa, including oysters (Farley 1969), mussels (*Mytilus trossulus*), soft-shell clams (*Mya arenaria*), common cockles (*Cerastoderma edule*), and golden carpet shell clams (*Polititapes aureus*). In cases where it has been determined to be a transmissible cancer, the lineages of contagious cancerous cells usually arise independently in each species (reviewed in Metzger et al.. 2016).

Common cytological diagnostic characteristics of neoplastic cells are large size, a high nucleus-to-cytoplasm ratio, rounded shape, non-adherent behavior in contrast to highly adherent hemocytes, and a prominent nucleus or several nuclei (Elston 1992; Barber et al. 2004; Carballal et al. 2013; Alderman et al. 2017; Dairain et al. 2020). There are several methods commonly used to diagnose DN in bivalves. Fluorescence-activated flow cytometry (FACS) is an established diagnostic technique to detect low levels of DN (<12% of total cells in hemolymph) as DN cells are usually polyploid, which can be detected with DNA-staining dyes (Mirella da Silva et al. 2005; Smolarz et al. 2005). Histological analyses of paraffin-embedded tissue sections from dissected animals or cytospin-generated hemolymph preparations can reliably detect neoplastic cells in advanced stages of disease (Carballal et al. 2013; Carballal et al. 2015; Dairain et al. 2020; Garcia-Souto et al. 2022). However, because neoplastic cells are often observed mainly in the hemolymph except in late-stage disease when other tissues are infiltrated by neoplastic cells, tissue sections can be of limited use as an early diagnostic tool in some systems (Barber et al. 2004).

The pathogenicity and prevalence of DN and its potential impacts on natural bivalve populations is not well understood. However, wide-scale mortality events due to DN outbreaks have been observed in mussels (*Mytilus* sp.) in the United States (Bower 1989, Barber 2004, Galimeny and Sunila 2008), soft-shell clams on the Atlantic coast of North America (Brown 1977, Barber 1990, Farley 1991, Muttray 2012), Baltic tellin (Dairain et al. 2020), and the common cockle in Europe (Le Grand et al. 2010). While reports of remission have been observed, infection is most often lethal (Elston et al. 1988, Barber 1990; Dairain et al. 2020). In the Pacific Northwest of the United States, a variety of bivalve species, some commercially important, are known to be susceptible to neoplasia (House and Elston 2006).

Through recent routine health screening of basket cockles (*Clinocardium nuttallii*) in Puget Sound, WA (USA), we observed evidence of DN in local wild populations. Basket cockles, a common bivalve native to the PNW, are a long-time traditional food staple for tribal communities (Apeti et al. 2013) and popular among recreational harvesters. There is growing interest in basket cockle aquaculture and potentially population enhancement among some Coast Salish tribes and the commercial industry (Dunham et al. 2013a, Dunham et al. 2013b), though recent observations of DN at a local broodstock source population (R. Elston, personal comm.) limits the viability of these projects.

To test large numbers of individual adult basket cockles for use as broodstock, quick, minimally invasive, and inexpensive screening for DN was needed. Here we present a hemolymph smear method for rapid live-screening of cockles for DN, using small amounts of hemolymph and inexpensive, accessible cell staining materials. We compare this method with established histological DN diagnostic techniques and show that H&E staining of hemolymph smears can be used to reliably identify DN in basket cockles. We suggest that this technique may be applied for detection of hemolymph-borne diseases in other bivalve species.

## 2. MATERIALS AND METHODS

### 2.1 Animal collection

Adult *Clinocardium nuttallii* (Figure 1A) were collected from beaches in in Puget Sound, WA in 2019: Semiahmoo (SM) in mid-December and Agate Pass (Y) in late March 2019.

**FIGURE 1:**
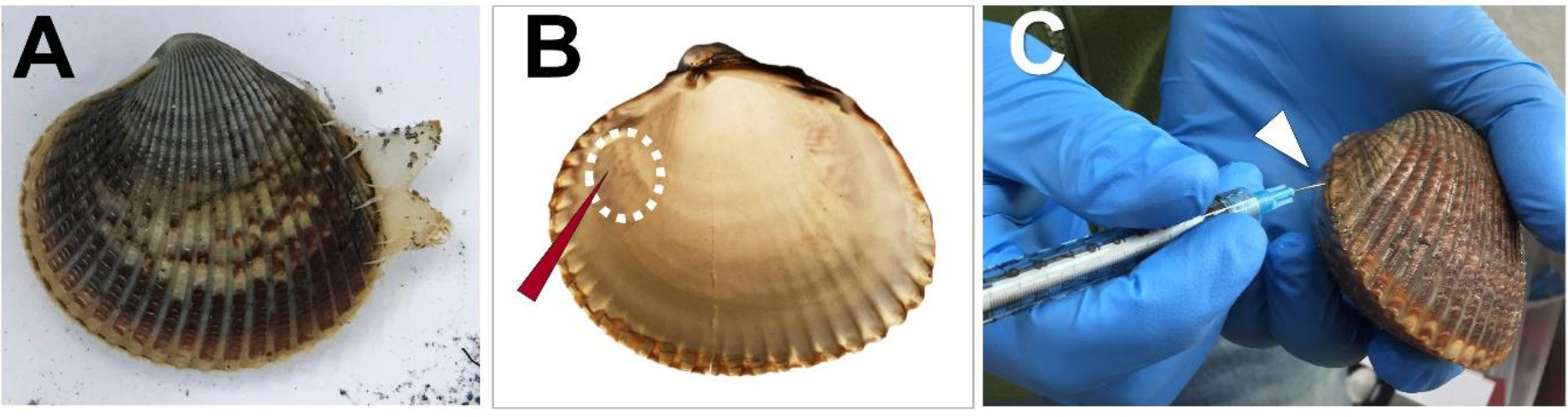
Demonstration of hemolymph draw in from *Clinocardium nuttallii*. A) Live basket cockle clam with siphons extended. B) Schematic representation of needle (red) position for drawing out hemolymph from the adductor muscle. C) Demonstration of minimally invasive hemolymph draw from a live cockle.

Previous routine disease assessments on cockles from this region showed a prevalence of neoplasia at roughly 10% (R. Elston, Washington Department of Fish and Wildlife, personal comm.). All cockle samples used in this study were collected in coordination with Suquamish Tribe shellfish biologists. Following collection, cockles were transported in coolers and processed for disease screening immediately upon arrival at the National Oceanographic and Atmospheric Administration’s Manchester, WA station.

### 2.2 Non-lethal hemolymph collection from C. nuttallii

To prevent stress-induced spawning or injury, we extracted a small volume of hemolymph from the adductor muscle using a 1 mL syringe and 26-gauge 1 inch (2.54 cm) needle (Figure 1B, C). This is done by gently handling an individual that had been resting undisturbed to minimize the risk that the valves are tightly shut, and inserting the needle while the valves are in a semi-open (relaxed) state. For this study, we collected a total of 300 μL of hemolymph from each individual cockle.

### 2.3 Sample Preparation for Histology

#### 2.3.1 Hemolymph Smear

To allow hemocytes to adhere to glass slides for processing, 100 μL of the collected hemolymph was applied to poly-L-lysine slides (Electron Microscopy Sciences, Hatfield, PA, USA) and allowed to rest for ten minutes at room temperature on a flat surface (Fig. 2A). A “blood smear” was performed by gently removing excess hemolymph using a coverslip (Fig. 2B,C). The slide air dried at room temperature on a flat surface. The adhered hemocytes were then fixed by submerging slides in 100% methanol for 15 minutes. The slides were removed from the methanol and air dried at room temperature. The slides were then in a covered container to prevent dust debris from attaching to the slides. Methanol-fixed hemolymph slides can then be stored at room temperature indefinitely or used immediately for histology.

**FIGURE 2:**
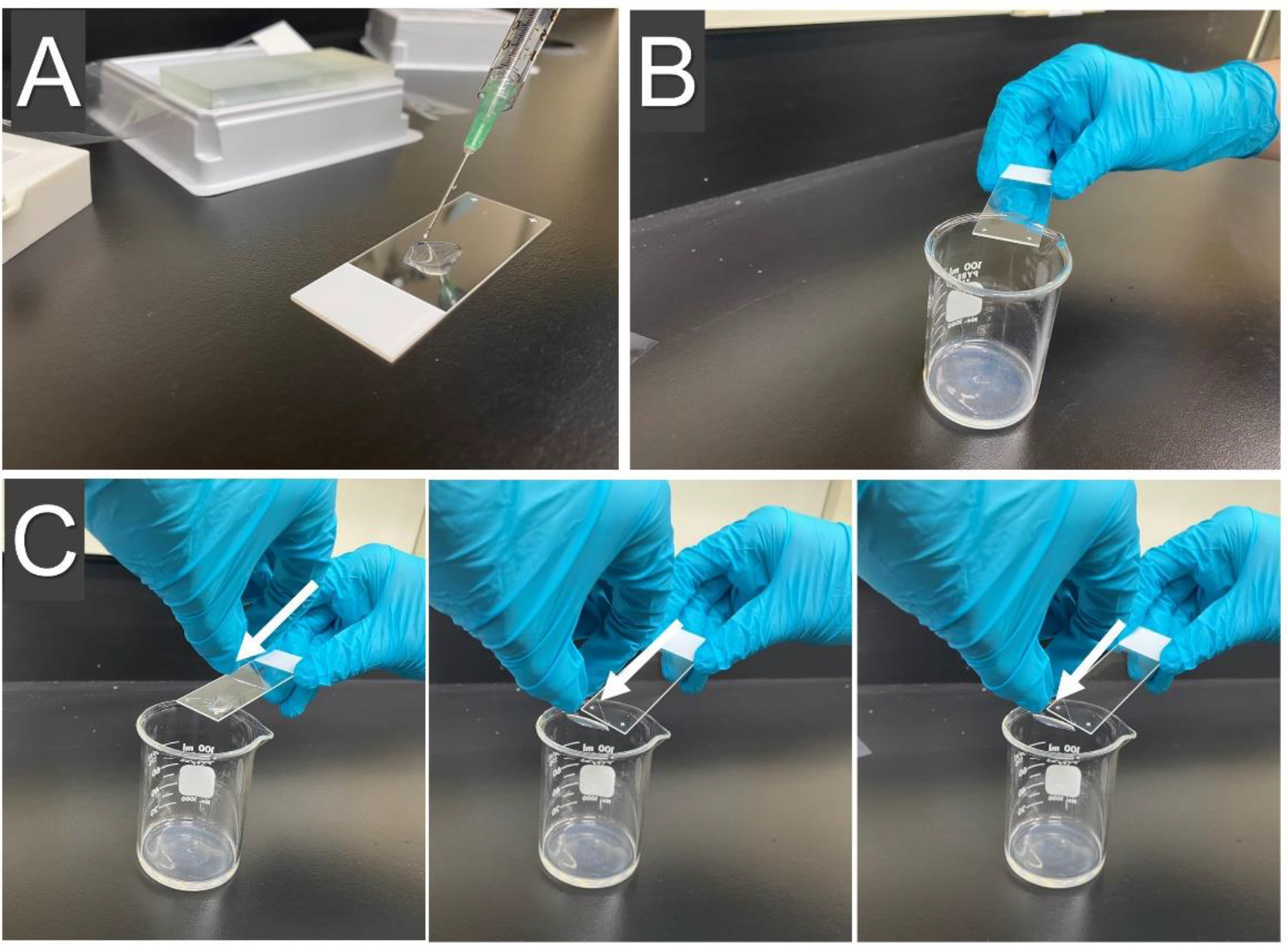
Demonstration of a hemolymph smear. A) Approximately 100 μL of hemolymph are applied to a poly-L-lysine-treated slide and allowed to rest 10 min on a flat surface. B) Position the slide over a waste container to collect excess hemolymph. C) Use a square coverslip to remove excess hemolymph by gently moving the coverslip across the slide surface from one end to the other, taking care not to make direct contact with the slide.

#### 2.3.2 Hemolymph Smear Hematoxylin and Eosin Staining

Note:All H&E reagents acquired from Leica, USA

1. Hydrate gradually in distilled water (70% EtOH → 50% EtOH → 25% EtOH) so fixed cells are not osmotically shocked and damaged.
2. Apply adequate Hematoxylin, Mayer’s (Lillie’s Modification) to completely cover smears on slides and incubate for 5 mins.
3. Rinse slide in two changes of distilled water to remove excess stain.
4. Apply adequate Bluing Reagent to completely cover the sample and incubate for 10-15 secs.
5. Rinse slides in two changes of distilled water.
6. Dip slides in 95-100% alcohol and blot excess ethanol off the edges of slides.
7. Apply adequate Eosin Y Solution (Modified Alcoholic) to completely cover the sample and incubate for 2-3 mins.
8. Rinse slide in 95% alcohol for 1 min.
9. Rinse slide in 100% alcohol for 1 min. Repeat twice more.
10. Dehydrate slides in three changes of ethanol (95% → 95% → 100%).
11. Clear slides in three washes of Hemo-De (d-Limonene) Xylenes substitute (Electron Microscopy Services) for one minute per wash.
12. Mount in synthetic resin, such as Permount (Fisher Scientific).

#### 2.3.3 Fixation, Embedding, and Sectioning of Whole Tissues

1. Excise thin section (∼2 mm) of tissue from gonad, stomach, foot, and gill using a scalpel.
2. Place in a tissue cassette, filling no more than 1/2 - 2/3 of cassette volume.
3. Transfer tissue cassettes to Davidson’s Solution (For 1L: 111 mL acetic acid, 320 mL 99% + ethanol, 222 mL 10% phosphate buffered formaldehyde solution, 367 mL de-ionized water) for 24-48hr. Note: Make sure volume of fixative is 10X volume of tissue.
4. Remove tissue cassettes from fixative and transfer to 70% ethanol. Cassettes can be stored at room temperature indefinitely.

## 3. RESULTS

### 3.1. Identification of DN in C. nuttallii through hemolymph analysis and histological methods

After DN was detected in cockle populations from Puget Sound, WA, based on standard, lethal histological methods (R. Elston, personal comm.), we developed the hemolymph smear protocol to non-invasively examine hemolymph morphology. When we collected cockles from Agate Pass, WA, we used a phase-contrast light microscope to observe that live hemocytes taken from non-neoplastic cockles exhibit diverse cell morphologies, with most cells being adherent and displaying pseudopodia, as has been observed in other bivalves (Fig. 3A; Carella et al. 2017). In contrast, live neoplastic cells do not readily adhere to tissue culture dishes and are rounded in shape, with an abnormally high density of cells in the hemolymph during advanced disease states compared to non-neoplastic individuals (Fig. 3B).

**FIGURE 3:**
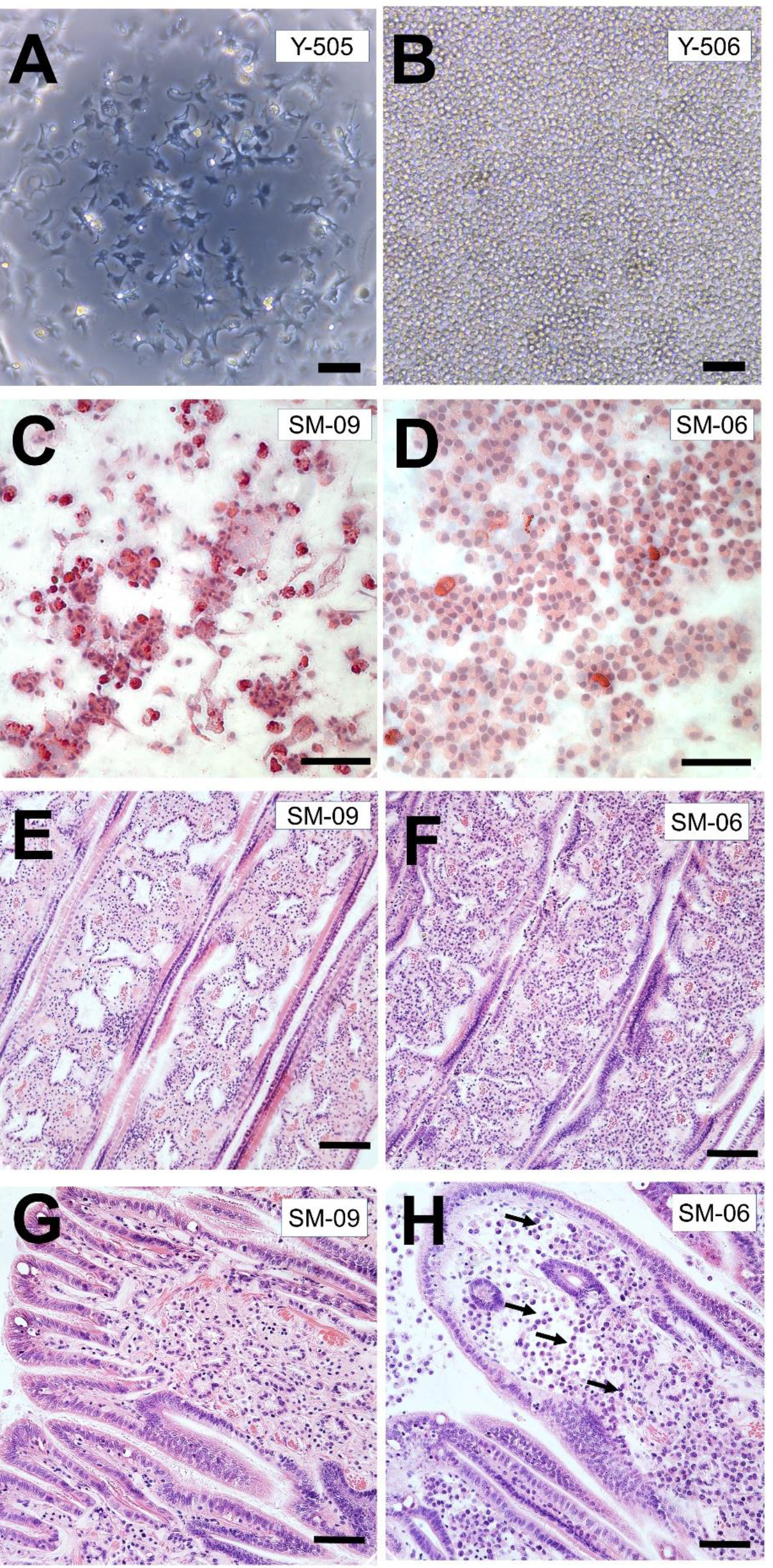
A) Live hemocytes in a non-neoplastic cockle sampled from Agate Pass, WA (sample Y-505). B) Mixture of live non-neoplastic hemocytes and neoplastic cells (white arrowheads) in an individual infected with DN sampled from Agate Pass, WA (sample Y-506). C) H&E stained hemocytes from a non-neoplastic individual collected from Semiahmoo, WA (sample SM-09). D) Cells stained by H&E displaying the distinctive rounded morphology and enlarged nuclei of neoplastic cells in a neoplastic individual collected from Semiahmoo, WA (sample SM-06). E-F) Hematoxylin and eosin histology of *C. nuttallii* gill tissue sections from animals collected from Semiahmoo, WA. E) Non-neoplastic gill tissue displaying normal tissue structure. F) Gill lamella of a neoplastic individual showing disorganized gross tissue structure and broad infiltration of neoplastic cells. G) High magnification of healthy gill tissue. H) High magnification of gill lamella from an animal with advanced DN. Arrows point to neoplastic cells. A-D: scale bar 50 μm. E, G: scale bar 200μm. F, H: scale bar 100μm.

When analyzing fixed cells from individuals from Semiahmoo using the hemolymph smear method, we observed non-neoplastic hemocytes displaying an array of cell morphologies and sizes (Fig. 3C), while cells taken from the hemolymph of an individual cockle with advanced neoplasia display a uniform size and round shape with enlarged nuclei, typical of the morphology of neoplastic cells (Fig. 3D; Barbet 2004). Standard histological analysis of cockle tissues confirms the diagnosis of DN, as determined by hemolymph analysis. Gill tissue from a non-neoplastic cockle shows normal tissue and cell morphology and cell density (Fig. 3E, 3G; compare to Fig. 3C). Conversely, a cockle with neoplasia displays gill tissue with abnormally high cell density and visible round cells with large nuclei (Fig. 3F, 3H; compare to Fig. 3D).

### 3.2 Validation of hemolymph smear protocol to detect disseminated neoplasia in basket cockles from Semiahmoo, WA, by direct comparison to histology

We screened individual cockles collected from Semiahmoo, WA (N=29) and identified neoplasia in two animals using the hemolymph smear method (6.7%). Methanol-fixed hemolymph smears show striking differences in cellular morphology between non-neoplastic and neoplastic individuals (Supplemental Fig. 1). To directly compare the accuracy of DN detection using the hemolymph smear method to tissue section histology, the same animals sampled for hemolymph smear staining were subsequently paraffin-embedded, sectioned, and H&E stained. Histological analysis of tissue sections confirmed the presence of neoplastic cells in the gill tissue in the same animals (SM-06, SM-26; Supplemental Figure 1), and no evidence of neoplasia was detected in the other 27 animals diagnosed as non-neoplastic using the hemolymph smear. These direct comparisons of the hemolymph smear method of DN detection to tissue histology using samples taken from the same individuals shows that non-invasive H&E staining of hemolymph smears is sufficient to accurately diagnose DN in *C. nuttallii*.

## 4. DISCUSSION

Disseminated neoplasia can be diagnosed using tissue histology, hemocytology, or flow cytometry, including fluorescence-activated flow cytometry (Carballal et al. 2013; Alderman et al. 2017). However, few research efforts have focused on large-scale screening methods for the purpose of surveying wild populations non-lethally. Rapid, inexpensive techniques for diagnosis of disseminated neoplasia in bivalves could be applied to disease ecology, fisheries management, population restoration efforts, and aquaculture.

Although considered common throughout their coastal Northeast Pacific Ocean range, the basket cockle, *C. nuttallii,* are relatively patchy in occurrence, making it difficult to monitor populations through established intertidal bivalve surveys employed by tribal and state fisheries. Available data suggest that *C. nuttallii* population dynamics are highly stochastic with temporally and spatially asynchronous population “booms” and “busts” (Barber et al.. 2019). Anecdotal reports of large recruitment and mortality events are commonly observed (B. Blake personal comm.); however, the factors regulating these population dynamics remain largely unknown.

The presence of DN in Puget Sound *C. nuttallii* populations compelled development of rapid, live sampling diagnostic methods to determine whether the disease was present in individual cockles intended for brood stock. Tissue histology is considered a standard approach for diagnosing DN in bivalves; it was therefore critical that we compared the accuracy of DN detection via the live-collected hemolymph smear method to classic tissue section histology (Barber 2004; Carballal et al. 2013; Carella et al. 2017). Examination of embedded and sectioned whole tissues compared to hemolymph smears from the same individuals show that the two techniques yield identical diagnostic results within the sample tested (Supplemental Figure 1). We posit that the minimally invasive hemolymph sampling and slide smear methods utilized in this study can robustly detect DN in cockle clams, equivalent to classical lethal whole-tissue sampling and histology methods.

Hemolymph smears are advantageous for multiple reasons including cost, speed, and technical logistics; they do not require access to specialized flow cytometry equipment, significantly reducing costs associated with disease diagnosis. In addition, hemolymph smears are relatively simple to carry out, and neoplastic cells are easy to discern via visual diagnosis. Lastly, this protocol does not require lethal sampling, allowing for repeated sampling of the same individual for research or diagnosis of animals destined for use as broodstock or captive display. This combination of sample collection and diagnostic tools has the potential to be generally applied to other taxa allowing for rapid, minimally invasive, and inexpensive diagnosis of DN in bivalves.

## ACKNOWLEDGEMENTS

The authors are grateful to Ralph Elston for technical support; Paul Hershberger and USGS Disease Ecology Laboratory, Marrowstone, WA; Suquamish Tribe Fisheries Department; Washington Department of Fish and Wildlife. The authors are also grateful to Pamela Johnson and the Benaroya Research Institute’s Histology Core facility for technical support.

## FIGURE LEGENDS

**SUPPLEMENTAL FIGURE 1:**
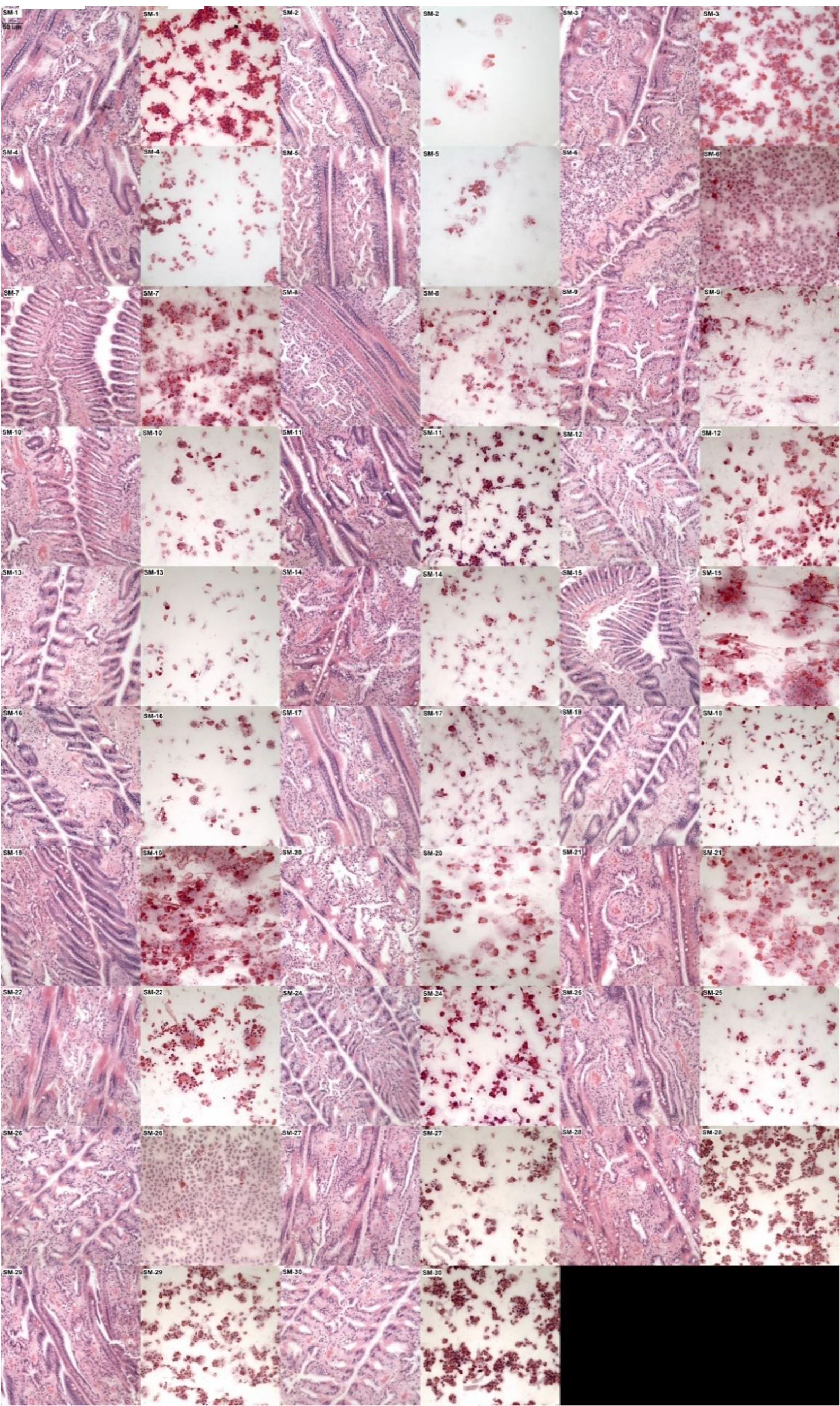
Hematoxylin and eosin histology of *C. nuttallii* gill tissue sections (left paired panels) compared to hemolymph smears (right paired panels). Neoplastic cells were detected in SM-06, and SM-26. All other samples appear to not have detectible neoplasia. Scale bar 50μm.

